## Supplementary Information for "African bush pigs exhibit porous species boundaries and appeared in Madagascar concurrently with human arrival"

### Supplementary Figures

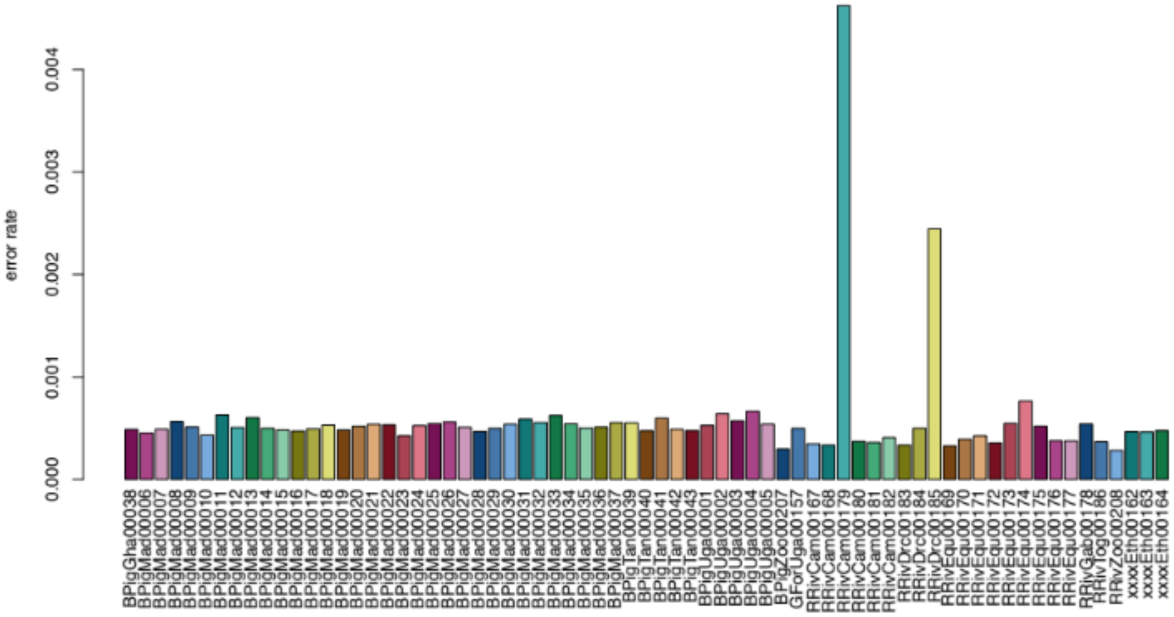

**Supplementary Figure S1. Estimated per-base sequencing error rates based on the ‘perfect individual‘ approach for samples mapped to the common warthog reference genome.**

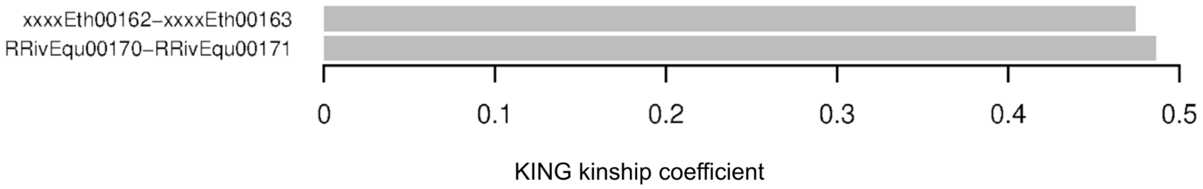

**Supplementary Figure S2. KING kinship coefficients for two Ethiopian and two Equatorial Guinean samples that were merged based on 2D-SFS.**

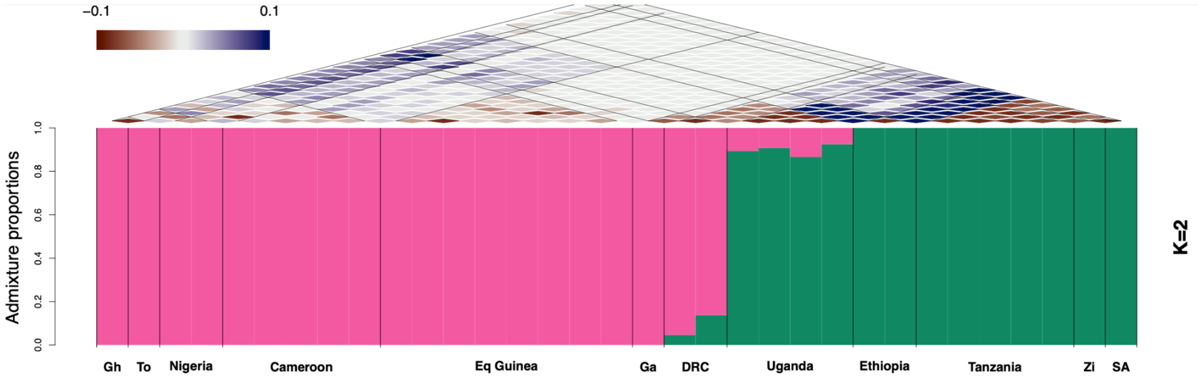

**Supplementary Figure S3. Inferred ancestry proportions for unrelated samples, excluding Madagascar (n = 33) using NGSadmix, assuming 2 (*K* = 2) ancestral populations.**

The barplot indicates admixture proportions for 2 ancestral populations, while triangles above indicate pairwise correlations of residuals as assessed by evalAdmix. Gh - Ghana, To - Togo, Ga - Gabon, DRC - Democratic Republic of Congo, Zi - Zimbabwe, SA - South Africa.

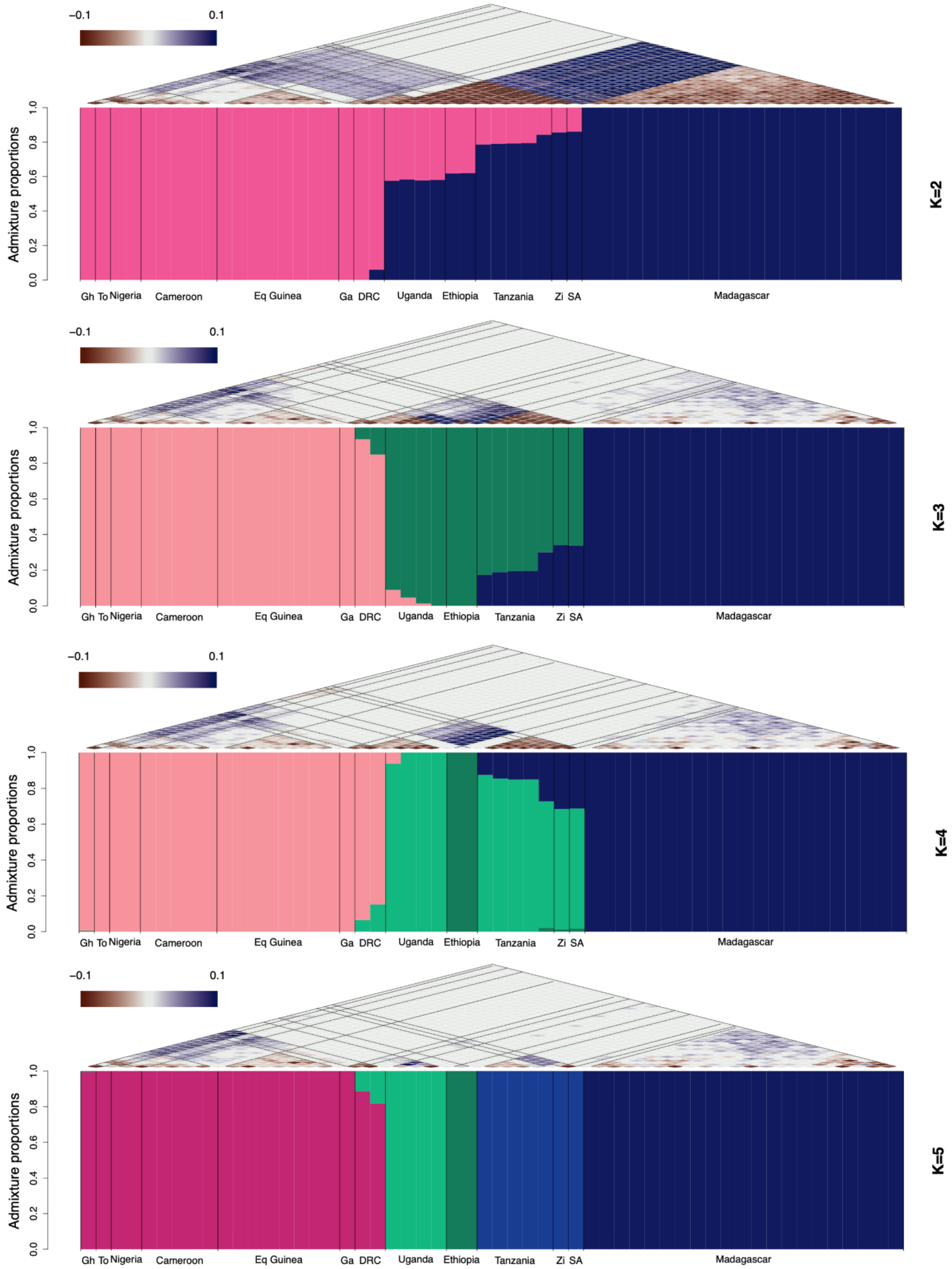

−0.1
0.1
Admixture proportions
0.0
0.2
0.4
0.6
0.8
1.0
Gh To Nigeria Cameroon Eq Guinea Ga DRC Uganda Ethiopia Tanzania Zi SA Madagascar
K=2
K=3
K=4
K=5

43

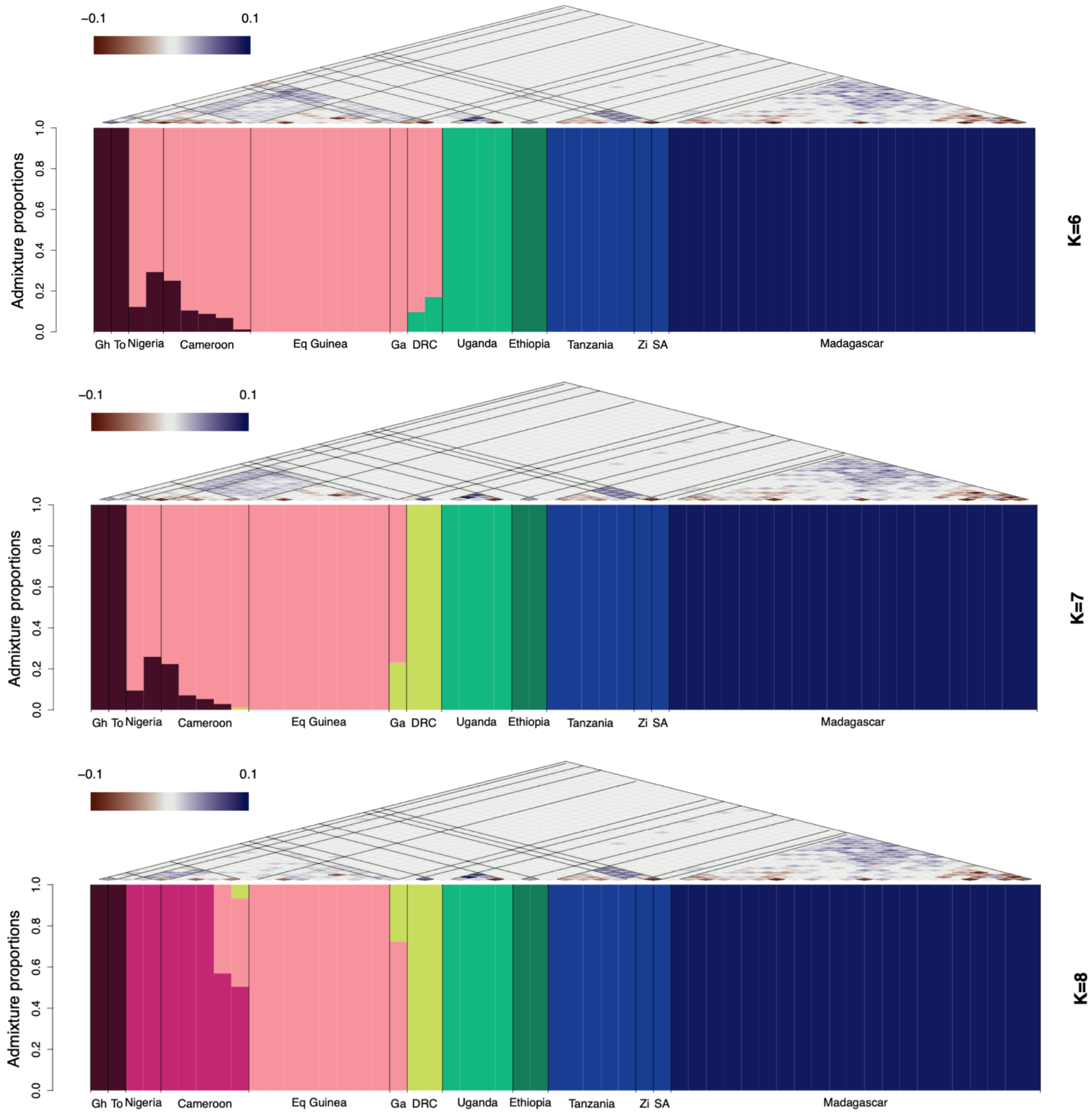

**Supplementary Figure S4. Inferred ancestry proportions for 54 unrelated samples using NGSadmix, assuming 2 (*K* = 2) to 8 (*K* = 8) ancestral populations.**

Barplots indicate admixture proportions for each number of assumed ancestral populations, while triangles above each barplot indicate pairwise correlations of residuals as assessed by evalAdmix. Gh - Ghana, To - Togo, Ga - Gabon, DRC - Democratic Republic of Congo, Zi - Zimbabwe, SA - South Africa.

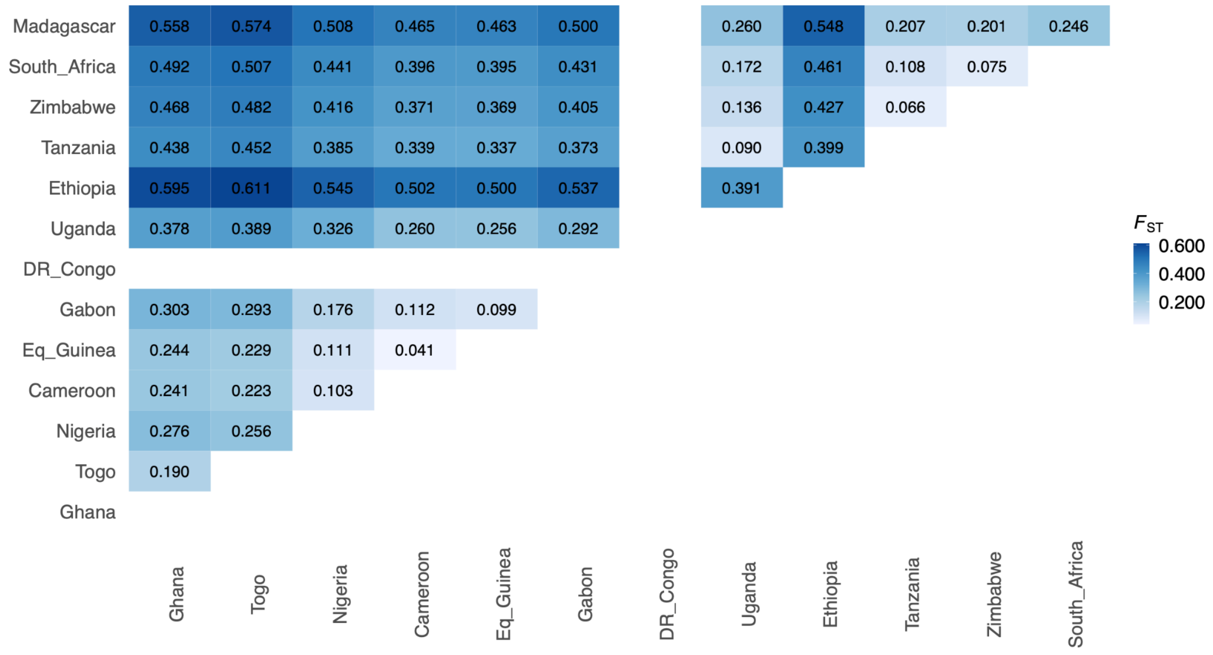

**Supplementary Figure S5. Hudson's $F_{ST}$ for 18 medium-high depth individuals.**

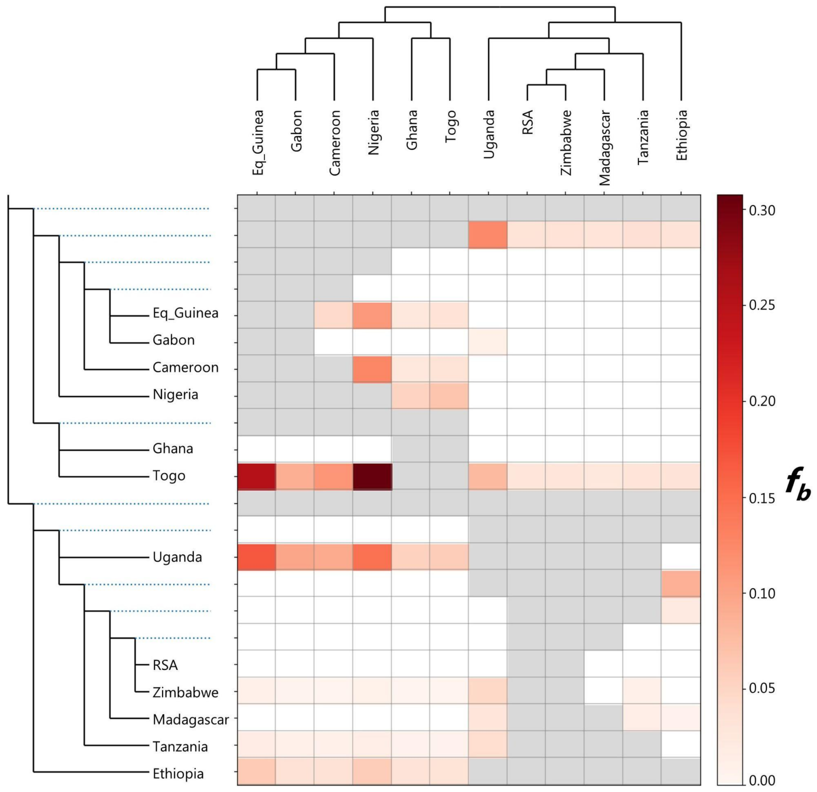

**Supplementary Figure S6. *f*-branch ($f_b$) statistics for all 13 populations.**
Heatmap indicates the extent of allele sharing between the corresponding branch of the species tree on the y axis, relative to its sister branch, and populations on the x-axis.

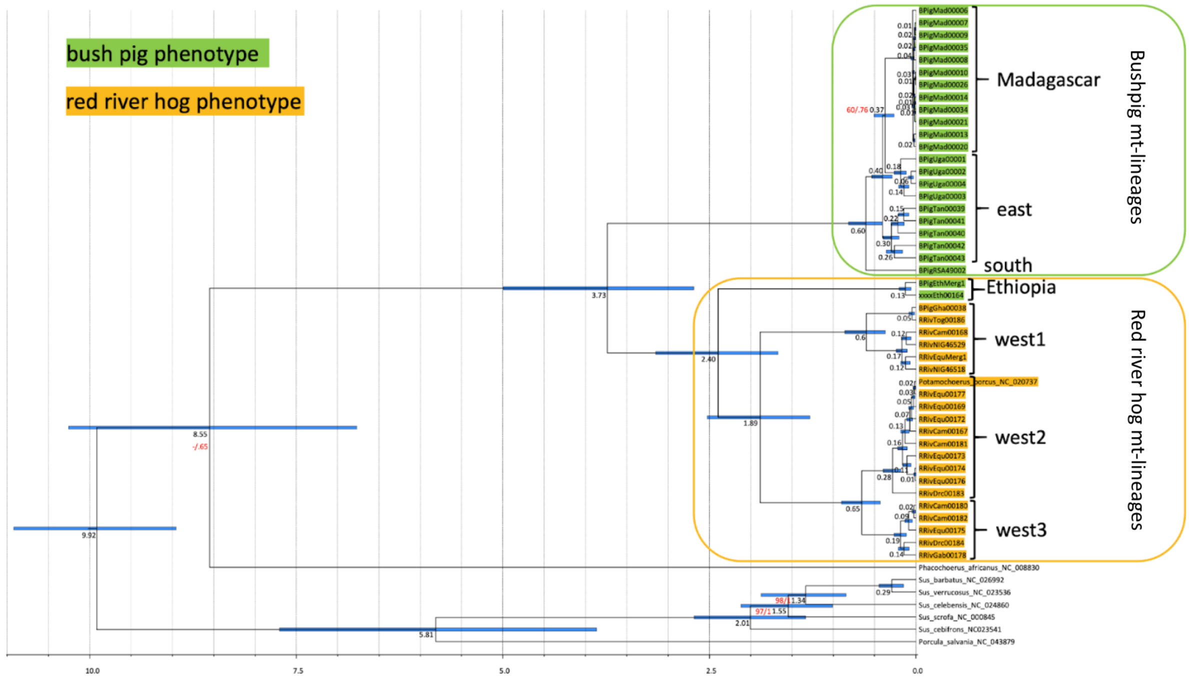

**Supplementary Figure S7. Mitochondrial phylogeny based on complete mitochondrial genomes using BEAST.**

### Supplementary Methods

#### Mitochondrial DNA phylogeny

Bioinformatic analyses were performed with the Geneious 2023.0.1 (https://www.geneious.com/) package. First, we trimmed and quality-filtered the reads with BBDuk 37.64 of the BBTools package (https://jgi.doe.gov/data-and-tools/bbtools/) and removed duplicate reads with Dedupe 37.64 (BBTools package), applying standard settings for both filtering steps. We then mapped the reads to the *Potamochoerus porcus* reference mitogenome (NC_020737) using the Geneious mapper with standard settings (medium-low sensitivity, fine-tuning with up to 5 iterations), annotated the mitogenome using the reference sequence as template, and checked the results by eye. Variant calling was verified with the find variations/SNPs tool in Geneious with standard settings (maximum variant $p$ value $10^{-6}$, minimum strand bias $p$ value $10^{-5}$ when exceeding 65% bias).

For phylogenetic reconstructions, we removed identical haplotypes (see Table S4) and added additional mitogenome sequences available in GenBank (*Potamochoerus porcus*: NC_020737, *Phacochoerus africanus*: NC:008830, *Porcula salvania*: NC_043879, *Sus scrofa*: NC_000845, *S. cebifrons*: NC_023541, *S. celebensis*: NC_026992, *S. barbatus*: NC_026992, S. *verrucosus*: NC_023536). We aligned the 52 sequences in the final alignment with Muscle 3.8.31 [1] in AliView 1.18 [2] and corrected the alignment by eye. We reconstructed a phylogenetic tree with the maximum-likelihood (ML) algorithm using IQ-TREE 2.2.0 [3]. Therefore, we treated the mitogenome as a single partition and applied the optimal substitution model (GTR + I + G) as determined with ModelFinder [4,5] in IQ-TREE under the Bayesian Information Criterion (BIC). To obtain node support for the ML analysis, we performed 10,000 ultrafast bootstrap (BS) replications [6].

We calculated divergence times in BEAST 2.6.7 [7] applying a relaxed lognormal clock model of lineage variation [8], a Birth Death tree prior and the best-fit model of sequence evolution as selected by ModelFinder. To calibrate the molecular clock, we set an age constraint on the node of the most recent common ancestor (MRCA) of Suinae with a mean of 10.0 million years ago (Mya) and a 95% highest posterior density of +/- 1.0 Mya [9,10]. We ran the analysis for 100 million generations with tree and parameter sampling setting in every 5,000 generations. To assess the adequacy of a 25% burn-in and convergence of all parameters, we inspected the trace of the parameters across generations using Tracer 1.6 (http://tree.bio.ed.ac.uk/software/tracer/). We combined sampling distributions of four independent replicates with LogCombiner and summarised trees with a 10% burn-in using TreeAnnotator (both programs are part of the BEAST package). We visualised all phylogenetic trees in FigTree 1.4.2 (http://tree.bio.ed.ac.uk/software/figtree/).
